# Targeting integrin αvβ3 by chimeric antigen receptor neural stem cell (CAR-NSC) therapy for stroke

**DOI:** 10.64898/2026.04.11.717950

**Authors:** Ruslan Rust, Rebecca Z. Weber, Nora H. Rentsch, Beatriz Achón Buil, Pardes Habib, Chantal Bodenmann, Kathrin J. Zürcher, Daniela Uhr, Debora Meier, Melanie Generali, Martina Zemke, Uwe Konietzko, Hirohide Saito, Simon P. Hoerstrup, Roger M. Nitsch, Christian Tackenberg

## Abstract

Stroke remains a leading cause of adult disability due to the brain’s limited regenerative capacity. Although stem cell therapies show favorable safety and feasibility profiles in early clinical trials, poor spatial retention and limited engagement with peri-infarct salvageable tissue constrain efficacy. Here, we engineered human induced pluripotent stem cell-derived neural stem cells (NSC) with a chimeric antigen receptor (CAR)-like architecture to enable targeted recognition of injury-associated cues. Specifically, cells were modified to express a membrane-anchored single chain variable fragment (scFv) targeting integrin αvβ3, a receptor selectively upregulated in peri-infarct vasculature after stroke. Engineered CAR-NSC retained progenitor identity and selectively bound recombinant integrin αvβ3 *in vitro*. Following focal transplantation into a photothrombotic stroke mouse model, CAR-NSC displayed broader dispersion within peri-infarct tissue and covered a greater proportion of the ischemic lesion compared to non-binding control-CAR-NSC. CAR-NSC grafts extended longer neurites that aligned more closely with the lesion border. In addition, CAR-NSC transplantation reduced microglial activation and was associated with increased vascular density and blood-brain barrier integrity in the peri-infarct zone. Together, these findings establish a CAR-like NSC strategy for stroke to direct the spatial distribution and tissue engagement of transplanted cells. Molecular targeting of injury-associated cues may improve the precision and regenerative efficacy of cell-based therapies for stroke and related neurological disorders.

## INTRODUCTION

Ischemic stroke occurs due to a sudden disruption of blood supply to the brain and results in permanent neurological deficits due to extensive neuronal loss and limited endogenous regeneration. It remains a leading cause of adult disability and mortality worldwide and is estimated to affect one out of four people over the age of 25 years in their lifetime ^1^. Although current reperfusion interventions, including thrombolysis and/or thrombectomy, improve survival, they have a narrow therapeutic time window and do not restore damaged neural circuits, leaving most patients with persistent functional impairments ^2^. Consequently, there is an urgent need for regenerative strategies that can promote functional recovery beyond the acute phase.

Stem cell-based therapy is an emerging approach to treat patients past the acute phase allowing functional recovery ^3^. Neural progenitor stem cells (NSC) derived from induced pluripotent stem cells (iPSCs) provide a promising cell source that is well-defined and scalable^4^. However, major limitations of these therapies are poor graft retention and a lack of spatial control over cell distribution within the complex, heterogeneous environment of the injury site ^5,6^. While NSC possess an intrinsic migratory capacity ^7–9^, they often fail to accumulate in the specific peri-infarct regions where they are most needed for structural and functional integration ^6^. The lack of precision to anchor cells to the salvageable tissue at the lesion border often results in grafts that remain sequestered within the necrotic core or disperse into healthy parenchyma.

To overcome these limitations, we applied genetic engineering to generate programmable therapeutic cells capable of recognizing and anchoring to injury-specific molecular cues. This approach draws inspiration from the clinical success of chimeric antigen receptor (CAR) T-cell therapies and the broader principles of synthetic biology, where synthetic receptors are used to couple environmental recognition with tailored cellular responses ^10,11^. Similar strategies have more recently been applied to monocytes and macrophages ^12,13^, and have also been extended to a brain-directed therapy using CAR-astrocytes targeting amyloid plaques in Alzheimer’s disease ^14^. Notably, CAR frameworks that decouple antigen recognition from intracellular effector signaling have also been explored, demonstrating that the extracellular targeting domain alone can be leveraged to spatially direct cell behavior.^15,16^

Here, by equipping NSC with engineered recognition domains (CAR-NSC), we hypothesized that we could transform them from passive grafts into spatially guided cell therapeutics following local administration. We engineered human iPSC-NSC as chimeric antigen receptor-like neural stem cells (CAR-NSC) by introducing a membrane-anchored single chain variable fragment (scFv) targeting integrin αvβ3. The construct employs the antigen-recognition principle of CAR architecture, with a non-signaling extracellular scFv designed to spatially retain grafted NSC within the αvβ3-enriched lesion border. αvβ3-CAR-NSC retained their neural identity and exhibited specific binding to their target ligand *in vitro.* Following focal transplantation into a photothrombotic stroke mouse model, αvβ3-CAR-NSC showed significantly enhanced dispersion and coverage of the ischemic stroke lesion compared to non-targeted control CAR-NSC. αvβ3-CAR-NSC further reduced microglial activation, promoted neurite outgrowth, and improved vascular density and blood-brain barrier integrity. Together, our findings establish antibody-guided cell surface engineering as a promising strategy to enhance the precision and regenerative potential of cell therapy for ischemic stroke.

## RESULTS

### Integrin αvβ3 is selectively upregulated in lesions after photothrombotic stroke

Integrin αvβ3 has been reported to be upregulated in the ischemic region after stroke ^17^. To verify αvβ3 integrin expression and protein abundance in our photothrombotic stroke mouse model, we first mapped expression of integrin αvβ3 in the photothrombotic stroke model across acute and subacute time points (1, 7, and 21 days post injury (dpi); **Fig. 1A**). Using immunostaining with anti-integrin αvβ3 antibody (clone LM609), we quantified αvβ3 signal in the injured hemisphere relative to the intact contralesional cortex and quantified the spatial distribution from the ischemic border zone (IBZ) **(Fig. 1B–E)**. Signal intensity of αvβ3 integrin increased over time and was spatially enriched in the infarct and peri-infarct tissue, with the highest signal detected approximately 250–400 μm from the IBZ (**Fig. 1C, D**). In contrast, no signal was observed in the intact hemisphere suggesting a specific upregulation of αvβ3 integrin in the stroke lesion (**Fig. 1C–E**). In the IBZ, αvβ3 integrin localized in close proximity to CD31+ blood vessels (**Fig. 1E**), consistent with previous observations ^18,19^.

**Figure 1:**
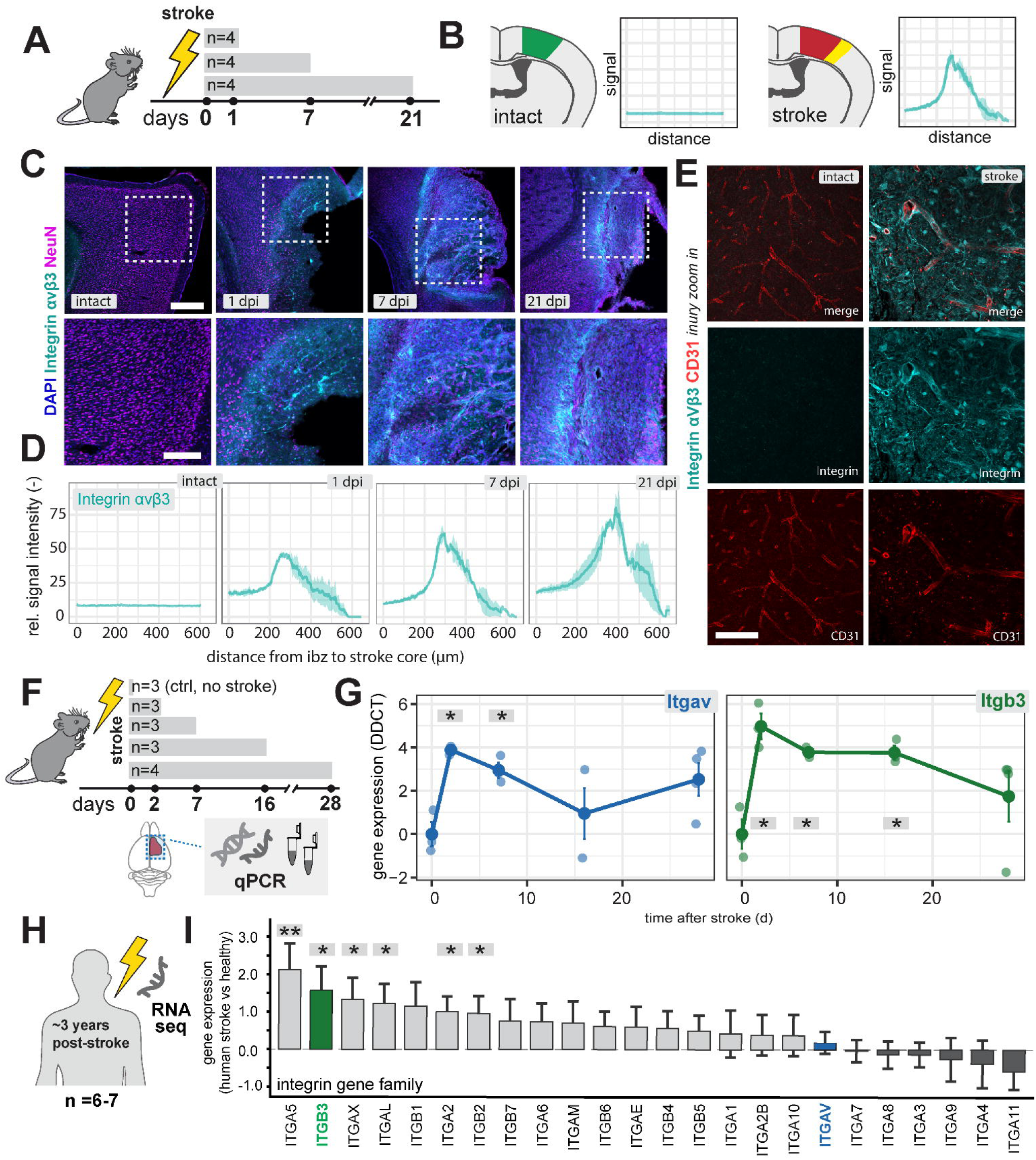
Integrin αvβ3 expression in mice after photothrombotic stroke. (A) Experimental timeline for photothrombotic stroke and tissue collection at 1, 7 and 21 days post injury (dpi). (B) Schematic of coronal brain sections illustrating regions analyzed, including peri-infarct and intact areas, with spatial distance measurement from stroke core. (C) Representative immunostaining of integrin αvβ3 (cyan), NeuN (magenta) and DAPI (blue) in intact cortex and peri-infarct tissue at 1, 7, and 21 dpi. Upper row: overview; lower row: higher magnification of boxed regions. (D) Quantification of integrin αvβ3 fluorescence signal as a function of distance from the stroke core in intact tissue and at 1 dpi, 7 dpi, and 21 dpi; n = 4. (E) High magnification images of integrin αvβ3 immunoreactivity with CD31 in peri-infarct tissue and intact cortex. (F) Experimental timeline and schematic for qPCR analysis of peri-infarct tissue at 2, 7, 16, and 28 dpi compared to non-stroke controls (n = 3, ctrl: no stroke). (G) Quantification of Itgav and Itgb3 mRNA levels in peri-infarct tissue, normalized to day 0; n = 3–4. (H) Schematic of reanalysis of publicly available RNA-seq data from post-mortem brain tissue of ischemic stroke patients (∼3 years post-stroke, n=7) and matched healthy controls (n=6) (GSE56267). (I) Differential expression of integrin gene family members in stroke versus healthy cortex. Bars represent log₂ fold change; error bars indicate ±1 standard error (DESeq2). ITGB3 (green) and ITGAV (blue) are highlighted. (G) Each dot represents one mouse. Scale bars: C (upper row), 100 μm; C (lower row), 50 μm; E, 20 μm. Data shown as mean ± SEM, *p < 0.05, one-way ANOVA with Dunnett’s test.

To corroborate these protein-level observations, we analyzed mRNA expression in peri-infarct tissue at 2, 7, 16, and 28 dpi and compared it to non-stroke brain tissue (**Fig. 1F**). Consistent with the histological data, both Itgav (αv integrin) and Itgb3 (β3 integrin) mRNA transcripts were significantly upregulated during the first week after stroke (**Fig. 1G**). Both Itgav and Itgb3 were sharply upregulated after stroke, with Itgav expression highest at 3-7 dpi, Itgb3 remained elevated through 16 dpi, supporting a sustained engagement of the αvβ3 integrin complex in the post-ischemic brain. To assess whether this upregulation extends to the human stroke, we reanalyzed publicly available RNA-seq data (GSE56267) from post-mortem brain tissue of stroke patients (median ∼3 years post-stroke; n=7) and matched controls (n=6)^20^. ITGB3 was among the most upregulated integrin family members, and both ITGB3 and ITGAV were elevated in stroke compared to control tissue (**Fig. 1H, I**), consistent with our observations in the mouse model.

Together, these data identify αvβ3 integrin as a temporally regulated and spatially enriched marker of the peri-infarct tissue in mouse and human stroke, motivating its use as a targeting cue for engineered CAR-NSC grafts.

### CAR-NSC preserve progenitor identity and enable specific αvβ3 integrin ligand binding *in vitro*

We differentiated human iPSCs into NSC using a previously established transgene- and xeno-free protocol ^4^. NSC stably expressed canonical stem cell and progenitor marker Nestin and lacked expression of the pluripotency iPSC-marker Nanog (**Fig. 2A**). Upon growth factor withdrawal, NSC spontaneously differentiated into neurons and astrocytes, confirming multilineage neural potential (**Fig. 2B**). To reduce the risk of iPSC residuals, cells were further purified using RNA switch (**Suppl. Fig. 1a**) ^4,21^, without altering NSC marker expression (**Suppl. Fig. 1b**).

**Figure 2:**
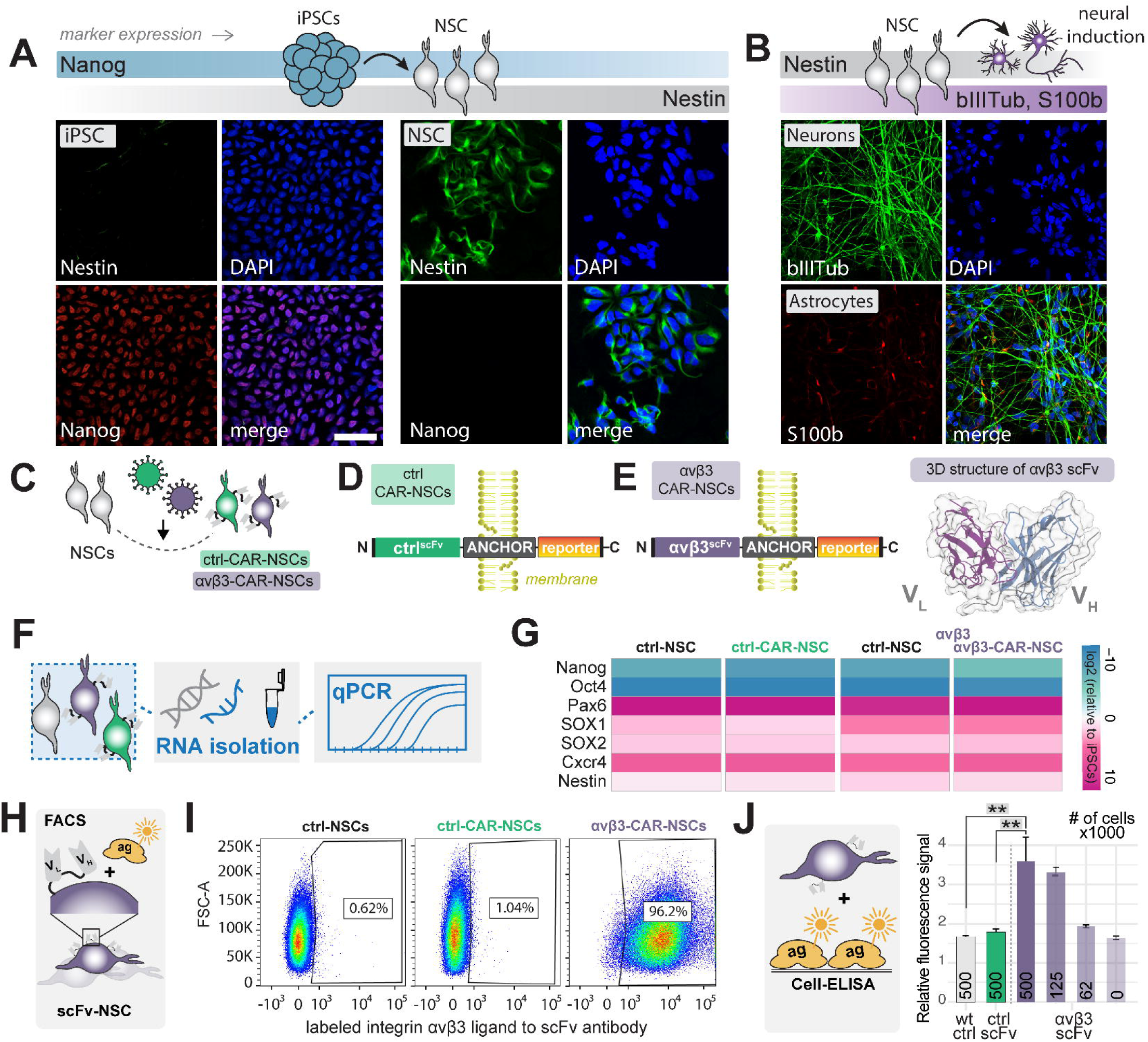
Generation and characterization of CAR-NSC expressing αvβ3 and control scFv constructs. (A) Left: Generation of human iPSC-derived NSC. Right: iPSCs and NSC (passage 7) stained for iPSC marker Nanog (red) and NSC marker Nestin (green), costained with Dapi (blue). Scale bar: 50 μm. (B) Left: Spontaneous neural differentiation of NSC. Right: Differentiated NSC at day 21 stained for neuronal marker βIII-Tubulin and S100β, a marker for astrocytes. (C) Schematic of lentiviral transduction of NSC to generate ctrl-CAR-NSC and αvβ3-CAR-NSC. (D) Construct design showing the scFv fused to a transmembrane anchor domain and intracellular reporter (rFluc or eGFP). (E) 3D structural prediction of the αvβ3-targeting scFv generated using AlphaFold3, showing VH and VL domains. (F) Schematic of RNA isolation and qPCR workflow for marker gene analysis. (G) Heatmap showing relative expression of iPSC markers (NANOG, OCT4) and NSC markers (PAX6, SOX1, SOX2, Cxcr4, NESTIN) in non-transduced NSC, ctrl-CAR-NSC, and αvβ3-CAR-NSC, analyzed by qPCR and displayed as fold change over iPSCs. (H) Schematic of flow cytometry assay for scFv-mediated ligand binding. (I) Representative flow cytometry dot plots of non-transduced NSC (left), ctrl-CAR-NSC (middle), and αvβ3-CAR-NSC (right) incubated with recombinant His-tagged αvβ3 integrin. Percentages indicate αvβ3-binding cells. (J) Cell-based ELISA (cELISA) showing Hoechst fluorescence intensity of cells bound to αvβ3 integrin-coated plates at indicated cell densities. Each bar represents the mean relative fluorescence signal of one well seeded with 500000 (wt, ctrl and αvβ3), 125000 (αvβ3), 63000 (αvβ3) or 0 cells. Statistical significance was assessed using Tukey’s HSD. *p < 0.05, **p < 0.01, ***p < 0.001.

To generate CAR-NSC with defined targeting specificity, we introduced a membrane-anchored single chain variable fragment (scFv) targeting integrin αvβ3 using a lentiviral construct under the EF-1α promoter (**Fig. 2C-E; Suppl. Fig. 2**). The scFv sequence was validated for binding to the integrin heterodimer and 3D structurally predicted using AlphaFold3 ^22^ (**Fig. 2E**).

As a non-binding control scFv, we used B12, an antibody specific for the HIV protein gp120^23^. The αvβ3 scFv was generated from the sequence of vitaxin, the humanized version of LM609 ^24^, which we used to verify integrin upregulation **(Fig. 1D, E)**. Vitaxin retains the properties of the LM609 antibody as determined by *in vitro* and *in vivo* characterization and has been proven safe in clinical applications ^24–26^. Both, control- and αvβ3-scFv have an N-terminal signal peptide and are C-terminally fused to a membrane anchor, for which the human EGF receptor transmembrane domain was used. To visualize NSC after transplantation via bioluminescence and fluorescence imaging ^27^, red firefly luciferase (rfluc) or eGFP were intracellularly fused to the membrane anchor as a reporter (**Fig. 2D-E, Suppl. Fig. 2**). Importantly, transduction with either control-scFv or αvβ3-targeting CAR construct did not measurably perturb the CAR-NSC state, as assessed by progenitor marker expression compared to non-transduced NSC (**Fig. 2G**).

We next tested whether CAR-NSC could specifically bind αvβ3 integrin *in vitro*. CAR-NSC were incubated with recombinant His-tagged αvβ3 integrin, and binding was quantified by flow cytometry using a fluorescently labeled anti-His antibody, with eGFP co-expression confirming successful CAR transduction (**Fig. 2H**). αvβ3-CAR-NSC showed robust ligand binding (96.2% positive), whereas ctrl-CAR-NSC and non-transduced wild-type NSC (ctrl-NSC) showed minimal binding (0.62% and 1.04%, respectively; **Fig. 2I**). To independently validate these findings, we performed a cell-based ELISA (cELISA) in which recombinant αvβ3 integrin was coated on plates and NSC binding was quantified by Hoechst fluorescence (**Fig. 2J**). Wild-type NSC and ctrl-CAR-NSC did not show fluorescence above background (**Fig. 2J**). In contrast, αvβ3-CAR-NSC showed significantly higher binding compared to all control groups (**p < 0.01). Notably, this binding was cell density-dependent, scaling with the number of seeded CAR-NSC (500,000, 125,000, and 62,000 cells; **Fig. 2J**).

As ctrl-CAR-NSC did not show binding to αvβ3 integrin, we verified the surface expression of the ctrl-CAR by protein L cELISA and flow cytometry. 96% of ctrl-CAR-NSC showed binding to protein L while only 1% of wild-type NSC were positive (**Suppl. Fig. 3A, B**). Furthermore, cELISA signal on protein L-coated plates was significantly higher for ctrl-CAR-NSC than for wild-type NSC (**Suppl. Fig. 3C**) indicating successful transduction and cell surface localization of the control-CAR.

Taken together, these data demonstrate that both ctrl-CAR and αvβ3-CAR are expressed on the NSC surface, but only αvβ3-CAR-NSC specifically bind αvβ3 integrin.

### Transplanted CAR-NSC show increased lesion coverage and neurite outgrowth

To determine whether αvβ3 recognition enhances graft positioning after stroke, we transplanted αvβ3-CAR-NSC or non-binding ctrl-CAR-NSC into the peri-infarct cortex one week after stroke (**Fig. 3A, B**). The timepoint was selected based on previous findings showing that delayed transplantation improves graft survival.^28^ A stroke group receiving sham transplantation (vehicle injection without cells) was included as a negative control. Stroke induction was validated by laser Doppler imaging (LDI), which confirmed consistent focal perfusion deficits (∼70% reduction in blood perfusion) across all experimental groups (**Fig. 3B, C**).

**Figure 3:**
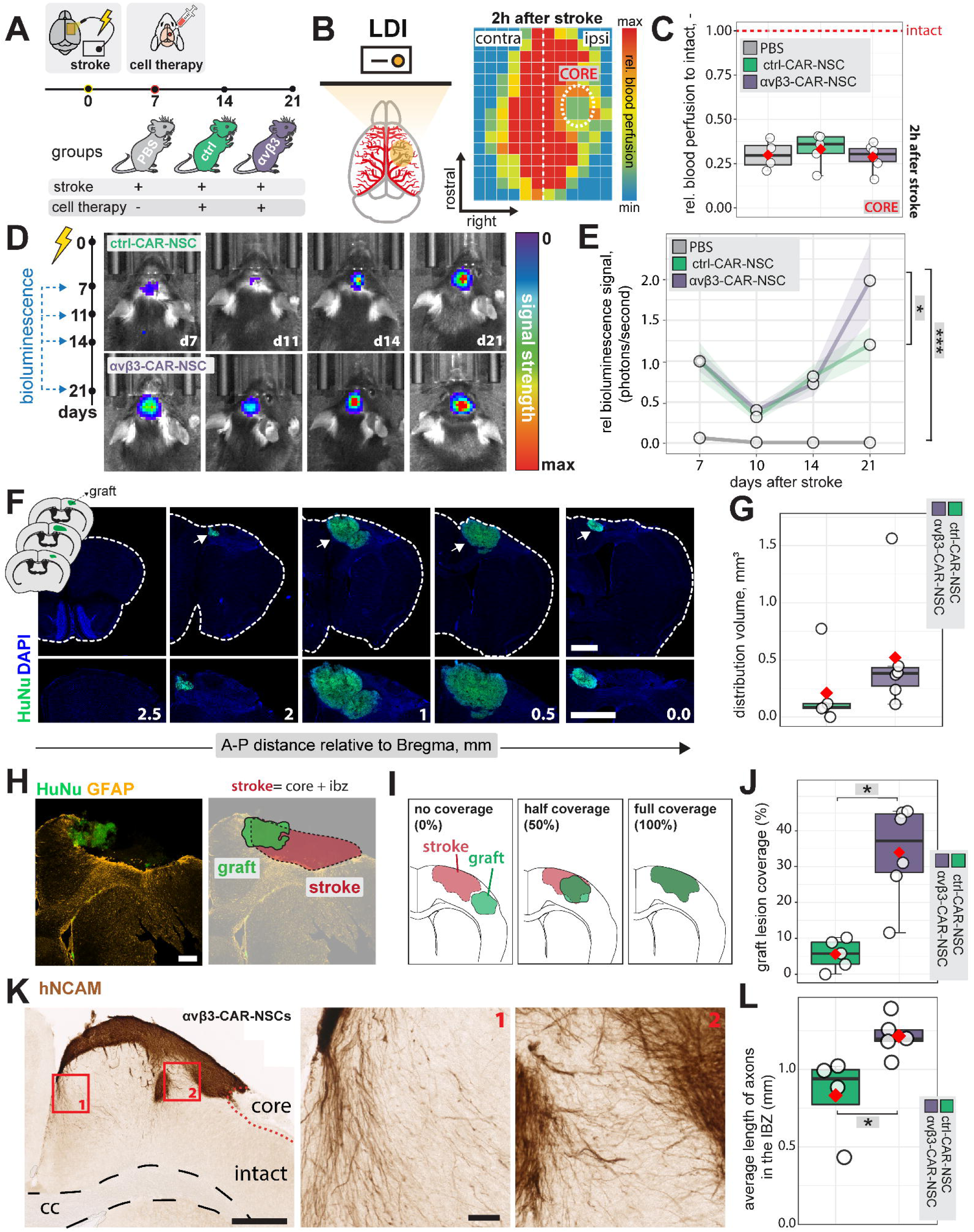
Graft survival, distribution, and lesion coverage of transplanted CAR-NSC. (A) Experimental design showing photothrombotic stroke followed by local transplantation of CAR-NSC at 7 dpi, with group allocation table. (B) Laser Doppler imaging (LDI) schematic and representative perfusion maps confirming focal reduction in cortical blood flow after stroke. (C) Quantification of relative blood perfusion, normalized to intact hemisphere, in PBS-receiving, ctrl-CAR-NSC-receiving and αvβ3-CAR-NSC-receiving groups; dashed red line indicates stroke threshold. (D) In vivo bioluminescence imaging (BLI) of ctrl-CAR-NSC and αvβ3-CAR-NSC groups over 14 days post transplantation, and 21 days post stroke respectively. (E) Quantification of bioluminescence signal (normalized photon flux) over time. (F) Representative coronal sections stained for Dapi (blue) and graft marker HuNu (green) at indicated anterior-posterior distances from bregma showing graft distribution (dashed outline). (G) Quantification of total graft distribution volume in ctrl-CAR-NSC and αvβ3-CAR-NSC groups. (H) Representative HuNu (green) and GFAP (orange) co-staining with corresponding 3D graft-lesion reconstruction schematic. (I) Schematic illustration of graft-lesion coverage categories: no coverage (0%), half coverage (50%), and full coverage (100%). (J) Quantification of graft lesion coverage (%). (K) Representative human-specific NCAM (hNCAM) immunostaining showing neurite outgrowth from αvβ3-CAR-NSC grafts in the peri-infarct cortex. Left: overview with intact cortex, stroke core (core), and corpus callosum (cc). Right: magnified views of boxed regions (1, 2). (L) Quantification of average neurite length in the ischemic border zone. Scale bars: F, 2 mm; H, 100 μm; K, 500 μm (left) and 50 μm (right). Data are shown as mean ± SEM; n = 6 (αvβ3-CAR-NSC), n = 5 (ctrl-CAR-NSC), n = 4 (PBS). Each dot represents one animal; boxes indicate 25th–75th percentiles with the mean shown. Line graphs plotted as mean ± SEM. In (E), statistical differences were assessed using one-way ANOVA followed by Bonferroni correction. In (J,L), statistical differences were assessed using an unpaired Wilcoxon–Mann–Whitney test due to non-normally distributed data. *p < 0.05, **p < 0.01, ***p < 0.001.

Longitudinal bioluminescence imaging (BLI) indicated survival of both graft types over 14 days post-transplantation (**Fig. 3D, E**). Signal intensity declined during the first week, suggesting early cell loss (as previously reported ^4,28,29^) followed by an increase of the bioluminescence signal from surviving grafts. Notably, at 14 days post transplantation, αvβ3-CAR-NSC exhibited higher bioluminescence signal compared to ctrl-CAR-NSC, indicating that scFv expression improved long-term viability (**Fig. 3E**).

We next assessed lesion volumes and graft distribution across the groups. GFAP staining was used to identify the ischemic lesion across anterior–posterior regions (**Suppl. Fig. 4A, B**). Quantitative volumetry revealed no reduction in lesion sizes in αvβ3-CAR-NSC–treated animals compared to controls (**Suppl. Fig. 4B,C**), indicating that the primary effect of αvβ3 targeting in this paradigm is not lesion reduction.

Given that integrin αvβ3 was found enriched at the ischemic border, we next asked whether αvβ3-scFv expression on CAR-NSC would influence graft distribution relative to the GFAP-defined lesion border. Human nuclei (HuNu) staining identified widespread graft presence within the peri-infarct and infarct cortex in both CAR-NSC groups (**Fig. 3F, G**). While both groups showed long-term engraftment, αvβ3-CAR-NSC tended to a 148% (p = 0.12) larger total distribution volume compared to ctrl-CAR-NSC (**Fig. 3G**). To quantify spatial graft-lesion relationships, we co-registered graft reconstructions with GFAP-defined stroke boundaries, which include the ischemic core and infarct border zone (**Fig. 3H**), and quantified graft-lesion coverage (**Fig. 3I**). Strikingly, αvβ3-CAR-NSC covered a significantly larger proportion of the ischemic lesion compared to ctrl-CAR-NSC (**Fig. 3J**), indicating preferential spatial alignment of αvβ3-CAR-NSC with the αvβ3-enriched peri-infarct border.

To assess whether αvβ3 targeting influences neurite outgrowth from grafted CAR-NSC, we analyzed projections from transplanted NSC using human-specific NCAM immunostaining in a separate cohort of animals at 28 days post-stroke (21 days post-transplantation), a timepoint at which more axonal sprouting is expected to occur (**Fig. 3K, Suppl. Fig. 5**). ctrl-CAR-NSC grafts displayed short, dense neurites that largely remained confined to the graft core and its immediate surroundings (**Suppl. Fig. 5**). In contrast, αvβ3-CAR-NSC extended longer and more radially oriented projections that penetrated deeper into the peri-infarct and intact tissue (**Fig. 3K**). Quantitative analysis showed that average axonal length within the ischemic border zone was significantly greater in mice receiving αvβ3-CAR-NSC compared to ctrl-CAR-NSC (p < 0.05, **Fig. 3L**).

Together, these results show that αvβ3-CAR-NSC achieve broader lesion coverage and enhanced neurite outgrowth in the peri-infarct region, supporting a role for αvβ3-mediated targeting in promoting graft integration in the post-stroke brain.

### CAR-NSC transplantation attenuates microglial activation independent of αvβ3 targeting

To investigate how CAR-NSC influence the post-stroke neuroinflammation, we assessed microglial/macrophage morphology in the peri-infarct region using Iba1 immunostaining at 21 days after stroke (**Fig. 4A**). Microglia in PBS-treated mice exhibit a morphology characterized by reduced branching and increased circularity, known to be associated with a reactive state.^30^ In contrast, both αvβ3-CAR-NSC and ctrl-CAR-NSC transplantation preserved a more ramified morphology resembling the contralateral hemisphere (**Fig. 4A-B**). Quantitative morphometric analyses (**Fig. 4C**) showed that transplantation of either CAR-NSC population significantly increased the perimeter while reducing circularity compared to PBS controls, indicating reduction in microglial activation in the ischemic border zone (**Fig. 4D-E**). No differences in branching index were detected across PBS, ctrl-CAR-NSC, and αvβ3-CAR-NSC groups (**Suppl. Fig. 6**) Notably, the immunomodulatory effect was comparable between αvβ3-CAR-NSC and ctrl-CAR-NSC, suggesting it was mediated by NSC transplantation per se, rather than αvβ3 targeting.

**Figure 4:**
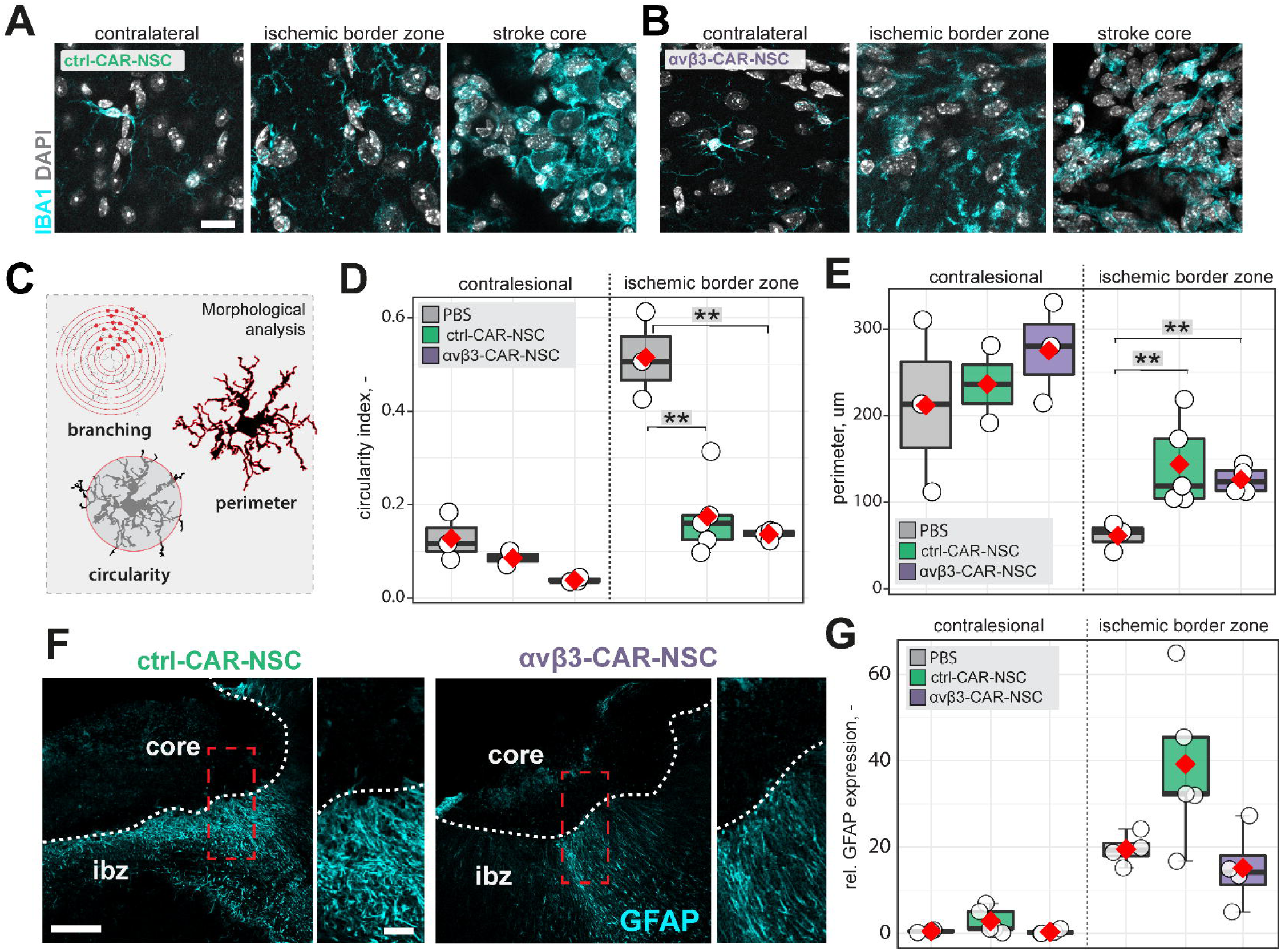
Microglial activation and astrocytic reactivity in PBS-, ctrl-CAR-NSC- and αvβ3-CAR-NSC-treated mice. (A,B) Representative images of Iba1 (cyan) and DAPI (grey) stained sections showing the contralesional cortex, ischemic border zone, and stroke core in ctrl-CAR-NSC- (A) and αvβ3-CAR-NSC-treated mice (B) at 21 dpi. Scale bar, 20 μm. (C) Schematic of morphological parameters quantified: circularity, and perimeter. (D) Quantification of microglial circularity. (E) Quantification of microglial perimeter. (F) Representative GFAP (cyan) staining of peri-infarct cortex in ctrl-CAR-NSC- (left) and αvβ3-CAR-NSC-treated mice (right), showing stroke core, ischemic border zone (IBZ), and GFAP-positive astrocytes. Scale bar, 50 μm (zoom: 25 µm) (G) Quantification of relative GFAP expression in contralesional cortex and ischemic border zone. Data are shown as mean distributions where the red dot represents the mean. Boxplots indicate the 25th to 75th percentiles. Each dot represents one animal. Statistical differences assessed by one-way ANOVA followed by Tukey’s HSD. *p < 0.05, **p < 0.01, ***p < 0.001.

We next examined astrocytic reactivity by quantifying GFAP expression in the peri-infarct cortex (**Fig. 4F**). Stroke induced strong GFAP^+^ astrogliosis in the peri-infarct region, as expected. However, GFAP^+^ expression levels were comparable between CAR-NSC-treated and PBS-treated animals (**Fig. 4G**), similar to previous findings.^29^

Together, these results indicate that CAR-NSC transplantation reduces microglial activation after stroke independent of αvβ3 targeting, while astrocytic reactivity remains largely unaffected.

### CAR-NSC increase vascular repair and BBB integrity in the peri-infarct zone

Prior studies have shown that NSC transplantation can promote vascular remodeling after stroke.^29,31,32^ To determine whether αvβ3-mediated targeting enhances vascular repair, we quantified vessel density, length, and branching in the ischemic border zone using CD31 immunostaining and automated morphometric analysis (**Fig. 5A-D**). Vascular parameters were assessed in defined regions of interest spanning the ischemic core, ibz, and intact tissue (**Fig. 5C**). PBS control mice showed a marked reduction in all vascular parameters compared to the contralateral hemisphere after stroke (dashed line; **Fig. 5D**). αvβ3-CAR-NSC significantly increased vascular area fraction compared to both PBS controls and ctrl-CAR-NSC (***p < 0.001 and *p < 0.05, respectively; **Fig. 5D**). Similarly, vessel length was significantly greater in αvβ3-CAR-NSC groups compared to PBS (*p < 0.001; **Fig. 5D**). No significant differences were observed in vessel branching across groups (**Fig. 5D**).

**Figure 5:**
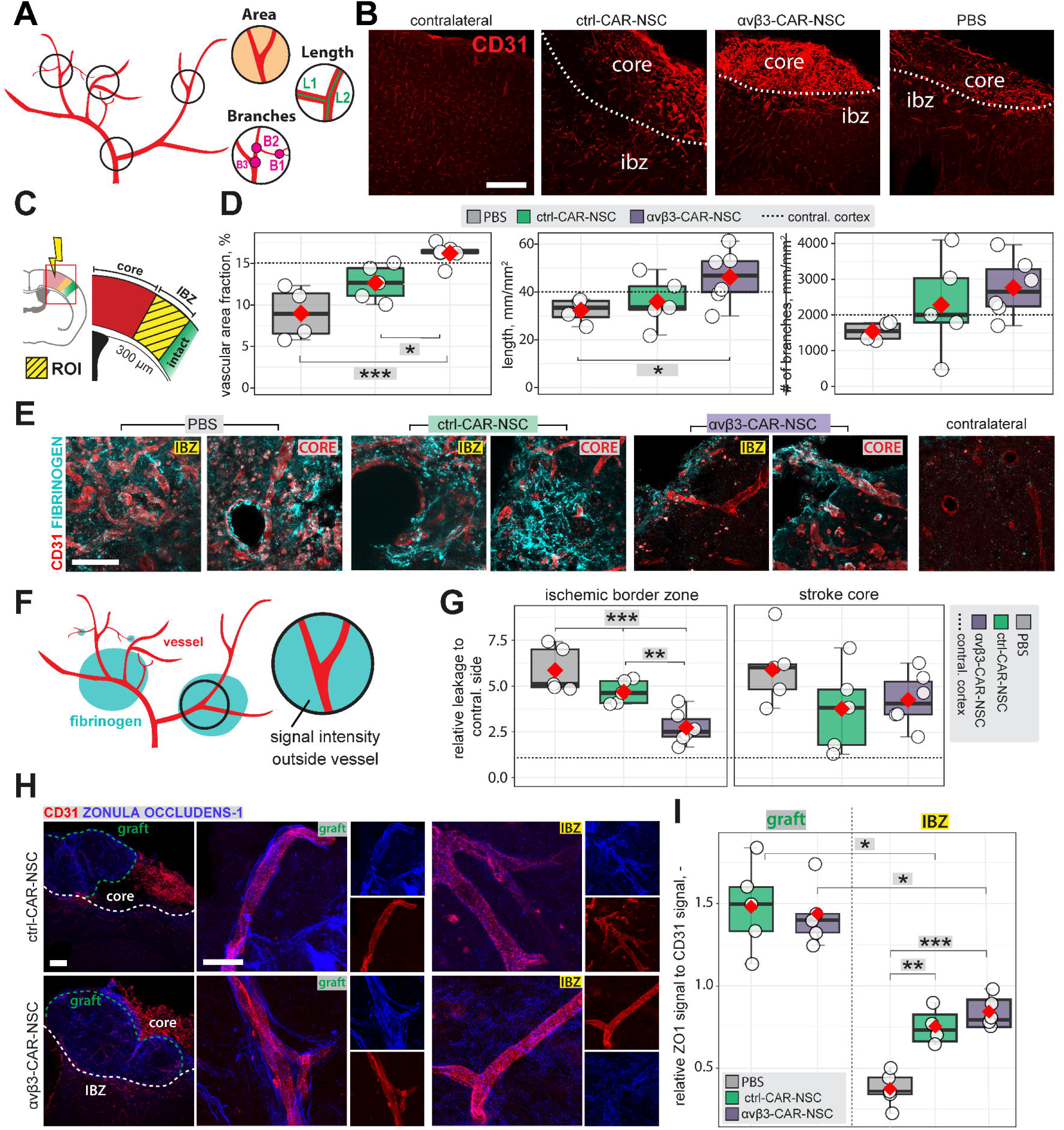
Vascular remodeling and BBB integrity in PBS-, ctrl-CAR-NSC- and αvβ3-CAR-NSC-treated mice. (A) Schematic of vascular parameters quantified: area fraction, vessel length, and branch points. (B) Representative CD31 (red) staining of microvessels in contralesional cortex and the ischemic border zone (IBZ) for PBS, ctrl-CAR-NSC and αvβ3-CAR-NSC groups, with stroke core and IBZ indicated. (C) Schematic of region-of-interest (ROI, yellow) placement spanning stroke core, IBZ, and peri-infarct zones. (D) Quantification of vascular area fraction, vessel length, and total branch points. Dashed line indicates contralateral cortex values. (E) Representative CD31 (red) and fibrinogen (cyan) immunostaining showing extravascular leakage in PBS-treated mice and reduced leakage in CAR-NSC-treated groups; contralateral cortex shown for reference. (F) Schematic of fibrinogen leakage quantification: signal intensity measured outside CD31-positive vessels. (G) Quantification of fibrinogen intensity in the ischemic border zone and stroke core. (H) CD31 (red) and ZO-1 (zonula occludens-1, blue) immunostaining in ctrl-CAR-NSC and αvβ3-CAR-NSC groups, showing stroke core and IBZ. (I) Quantification of the ZO-1/CD31 intensity ratio in graft-associated regions and the IBZ. Scale bars: B, 100 μm; E, 10 μm; H (left), 100 μm; H (right), 10 μm. Data are shown as mean distributions with the red dot indicating the mean. Boxplots indicate the 25th to 75th percentiles, and each dot represents one animal. Statistical differences were assessed using one-way ANOVA followed by Tukey’s HSD. *p < 0.05, **p < 0.01, ***p < 0.001.

To assess blood–brain barrier (BBB) integrity, we measured extravascular fibrinogen as a marker of leakage (**Fig. 5E, F**). PBS-treated mice displayed pronounced fibrinogen accumulation in the ischemic border zone (**Fig. 5G**). Both CAR-NSC groups showed reduced leakage compared to PBS, with αvβ3-CAR-NSC showing the greatest reduction, significantly lower than both PBS (***p < 0.001) and ctrl-CAR-NSC (**p < 0.01; **Fig. 5G**). No significant differences were observed in the stroke core (**Fig. 5G**).

To better understand how NSC transplantation contributed to BBB integrity, we looked at tight junction organization and analyzed ZO-1 expression relative to CD31+ vessels in graft-associated regions and the ischemic border zone (IBZ) (**Fig. 5H**). Quantification revealed significantly higher ZO-1/CD31 ratios in both ctrl-CAR-NSC and αvβ3-CAR-NSC treated mice compared to PBS controls (**Fig. 5I**). This increase was evident both in vessels directly adjacent to the graft and in the surrounding IBZ. Notably, no differences were detected between ctrl-and αvβ3-CAR-NSC groups, suggesting that the effect was mediated by NSC transplantation per se rather than αvβ3 targeting.

Together, these findings demonstrate that CAR-NSC transplantation promotes vascular remodeling in the peri-infarct region, as evidenced by increased vessel density and length. Notably, αvβ3-CAR-NSC showed superior BBB restoration, reducing fibrinogen leakage and enhancing tight junction organization in the ischemic border zone, consistent with their preferential spatial alignment along the αvβ3-enriched peri-infarct vasculature.

## DISCUSSION

Stem cell-based therapies for stroke have demonstrated feasibility and safety in early-phase clinical trials, yet their efficacy remains constrained by poor spatial control over graft distribution within the injured brain^33–36^. Here, we demonstrate that human iPSC-derived NSC can be engineered as CAR-like NSC to recognize injury-associated molecular cues and more effectively localize to peri-infarct tissue after stroke. We identified the αvβ3 integrin heterodimer, which is upregulated in the peri-infarct vasculature, as a suitable molecular target to guide NSC-based therapies. Expression of an αvβ3-targeting single chain variable fragment (scFv) on the NSC surface enabled selective binding to its ligand *in vitro* while preserving NSC identity. Following transplantation, αvβ3-CAR-NSC showed broader distribution and greater engagement with the peri-infarct tissue than non-binding ctrl-CAR-NSC. Together, these findings show that directing stem cells toward injury-defined molecular signatures improves the spatial positioning of transplanted cells, establishing a CAR-like strategy to enhance their therapeutic potential for stroke.

We observed selective upregulation of αvβ3 protein and transcript in peri-infarct tissue across acute and subacute time points, with peak enrichment at 250–400 μm from the ischemic border zone, a region considered most permissive for graft integration and trophic support ^37,38^. These findings are consistent with prior studies demonstrating αvβ3 induction in peri-infarct regions across rodent stroke models ^17,39–41^, as well as in non-human primates and human stroke patients ^42–44^. Histological analyses localized this increase to microvessels within ischemic brain tissue, linking αvβ3 expression to vascular remodelling after stroke ^18,19^. In addition, αvβ3 and its ligand osteopontin have been reported to accumulate within the glial scar, suggesting a broader role in extracellular matrix remodeling during post-stroke tissue reorganization ^18^. Together, these data support αvβ3 as a robust and conserved feature of the post-stroke brain and a promising molecular guidance cue for targeted stem cell-based therapies.

Notably, our time course analysis showed that both *Itgav* and *Itgb3* remained upregulated beyond the acute phase, suggesting sustained Itgb3 expression during post ischemic remodeling, consistent with observations in human stroke brain mRNA even at a median of three years post stroke (**Fig. 1H-I**). This pattern indicates that the αvβ3 targeting window extends into the subacute phase, a time frame that is clinically relevant for cell transplantation. Beyond stroke, αvβ3 is also upregulated in the ischemic myocardium after myocardial infarction ^45–47^. Accordingly, engineering iPSC-derived cardiomyocytes (iCMs) with an αvβ3-targeting CAR-like construct may enhance their retention within ischemic tissue and improve the efficacy of cardiac regenerative therapies, a possibility that warrants direct experimental testing.

Our findings also suggest that αvβ3-CAR-NSC occupy a larger graft volume and align more closely with the peri-infarct border, whereas non-binding control CAR-NSC remain more confined to the injection site. This improved spatial distribution is accompanied by longer and radially oriented neurites extending into peri-infarct tissue, possibly suggesting enhanced structural engagement with the host environment. The peri-infarct ECM is enriched in known αvβ3 ligands including fibronectin and osteopontin, which may provide a more permissive substrate for neurite extension in grafts that are better anchored within this niche ^18^. The mechanism by which αvβ3 binding enhances graft distribution likely involves adhesion-mediated retention along αvβ3-expressing peri-infarct vasculature, which may serve as a physical scaffold for cell migration and anchoring.

Unlike conventional CAR constructs used in T cell therapy, which include intracellular signaling domains (e.g., CD3ζ, 4-1BB) to activate effector functions upon antigen binding ^48,49^, our CAR-NSC design consists exclusively of a membrane-anchored scFv without intracellular signaling. This approach is conceptually related to non-signaling CARs^12,49^, which have been used e.g., to direct γδ T cells to tumor targets through antigen recognition alone ^50^, or to enhance T cell adhesion to stromal antigens via non-signaling antibody-binding scaffolds^51^. We therefore refer to our construct as CAR-like: it retains the chimeric antigen recognition architecture but relies on spatial retention within the αvβ3-enriched border zone rather than intracellular signal transduction.

The relevance of this finding is supported by earlier studies showing that transplanted neural stem cells can disperse widely within the brain and that functional recovery is greatest when grafts reside in peri-infarct regions rather than within the necrotic core, where the microenvironment is less permissive for integration and trophic support ^37,38^. Effective lesion coverage is therefore likely a key determinant of therapeutic efficacy, as closer graft–host contact have shown to promote synaptic remodeling, trophic signaling, and vascular interactions^37,38^. However, we did not directly assess synaptic integration of grafted neurons, and the functional significance of the observed neurite extension remains to be established.

Importantly, αvβ3 targeting did not reduce lesion volume, which is consistent with the relatively short observation window (21 days) and the known timeline of structural tissue remodeling after stroke. Lesion volume reduction is unlikely to be a primary readout of graft positioning effects at this time point, and its absence should not be interpreted as a lack of therapeutic engagement. Notably, this is in line with previous studies, in which NSC-treated animals showed improved gait and fine-motor recovery despite no significant reduction in lesion volume, underscoring that lesion size does not necessarily predict functional recovery after stroke ^28,29^. In the context of larger human stroke lesions, where distances between grafts and salvageable stroke tissue are substantially greater than in mice, targeted enrichment at the lesion border may become even more critical for achieving meaningful functional repair. Additionally, from a clinical perspective, intracerebral transplantation within the peri-infarct region carries the risk of damaging intact tissue, which makes direct placement in the peri-infarct cortex less desirable. Injecting cells into the stroke core and enabling them to migrate toward salvageable tissue may therefore represent a safer and more practical strategy. In this setting, CAR-NSC that preferentially accumulate at the lesion border could offer a distinct advantage over unmodified cell preparations. This becomes especially relevant for systemic blood-infusion delivery approaches, where homing cues are essential and integrin-based targeting could synergize with BBB shuttle strategies ^52^. Future studies will need to evaluate the efficacy of systemic delivery paradigms of engineered NSC.

Both αvβ3-CAR NSC and ctrl-CAR-NSC reduced microglial activation and improved blood brain barrier integrity, consistent with our previous findings ^28,29^. This effect was not enhanced by αvβ3 targeting, suggesting it reflects a general paracrine property of NSC transplantation rather than a consequence of improved spatial positioning. Notably, αvβ3-CAR-NSC also increased vascular density and vessel length, indicating that spatial distribution may enhance their paracrine influence on the post stroke vasculature ^31,32^. This dissociation, whereby BBB stabilization (fibrinogen, ZO-1) was a general NSC effect while increased vascular density was specific to αvβ3-targeted grafts, suggests that anchoring grafts closer to remodeling host vessels may selectively enhance proximity-dependent paracrine pro-angiogenic support and thereby promote vascular remodeling and BBB restoration. However, because the scFv derives from the function-modulating antibody vitaxin/LM609, direct modulation of host αvβ3 signaling on endothelial cells cannot be excluded as a contributing factor to the observed vascular effects.

Our study has the following limitations. First, although we have previously shown that Rag2⁻/⁻ mice recapitulate stroke pathology comparable to wildtype C57BL/6J mice^53^ and support long-term graft survival similar to clinically used immunosuppression strategies such as tacrolimus^53^, the use of immunodeficient models may still limit direct clinical translation, as immune responses to grafted cells could influence graft survival and therapeutic efficacy. Future studies could address this by employing engineered iPSC-NSC lines designed to evade immune rejection^54,55^ and enable allogeneic cell therapy approaches for stroke. Second, although grafted CAR-NSC differentiated into mature neurons and extended long range axonal projections, we did not directly assess their functional synaptic integration. Third, all transplantations were performed locally into the peri-infarct cortex; the efficacy of αvβ3-guided targeting under systemic delivery conditions, where homing cues are essential, remains untested. As a control, we used ctrl-CAR-NSC expressing an irrelevant scFv, which matches vector backbone, promoter, membrane anchor, and transduction history, thereby isolating the effect of αvβ3-specific binding. However, this design did not include unmodified NSC and thus does not control for potential effects of lentiviral transduction or CAR surface display per se on NSC. Finally, we did not perform behavioral analyses to determine whether αvβ3-CAR-NSC targeting translates into improved functional long-term recovery. Future work will be required to link enhanced graft targeting and tissue repair to defined behavioral outcomes and recovery mechanisms.

In summary, our study demonstrates that human iPSC-NSC can be engineered with a CAR-like architecture to recognize injury-associated molecular cues and more effectively localize to defined regions of the injured brain after stroke. This CAR-like guided engineering is modular, as the scFv targeting domain can be adapted to engage distinct injury-associated ligands and, if desired, coupled with intracellular signaling domains to elicit cell-autonomous responses. In principle, this strategy could be extended to other therapeutic cell types and disease contexts, providing a general framework for molecularly guided regenerative cell therapies.

## MATERIALS AND METHODS

### Experimental Design

This study was designed to evaluate the targeting and regenerative potential of human iPSC-derived neural progenitor stem cells (NSC) engineered to express a single-chain variable fragment (scFv) fused to a membrane anchor and luciferase or fluorophore reporter. The scFv construct was designed to specifically bind integrin αvβ3, which is known to be upregulated in the ischemic stroke area.

To investigate in vivo targeting and survival, immunodeficient Rag2⁻/⁻ mice were subjected to photothrombotic stroke induction and randomly assigned to one of three groups: (1) αvβ3 - scFv–expressing NSC transplantation (n = 6), (2) control NSC transplantation (n = 5), or (3) PBS injection (n = 4). Transplantations were performed locally into the stroke cavity seven days after stroke induction.

To monitor NSC survival, bioluminescence imaging (BLI) was performed on the day of transplantation, four days later, and weekly thereafter for a total of 14 days post-transplantation (21 days after stroke). After sacrifice, histological analyses were conducted to assess graft homing, lesion coverage, neurite outgrowth, and graft size. Additional analyses included quantification of stroke volume, microglial activation, vascular parameters, and astrogliosis. All animals included in the study are reported; no statistical outliers were excluded.

Both sexes were used and analyzed for this study; our statistical analyses did not reveal any significant sex-related differences in the measured parameters.

### Animals

All animal experiments were conducted at the Laboratory Animal Services Center (LASC) in Schlieren according to the local guidelines for animal experiments and were approved by the Veterinary Office in Zurich, Switzerland (license No: ZH110/2023, ZH209/2019). Adult male and female genetically immunodeficient *Rag2*^-/-^ mice (weight range: 20-30g) were employed for this study. Mice were maintained at the LASC in top-filter laboratory cages under standard housing conditions with controlled temperature and humidity, and a 12/12h light/dark cycle (light on from 6:00 a.m. until 6:00 p.m.). All mice were housed in groups of two to four per cage with ad libitum access to standard food pellets and water.

### NSC differentiation and culture

Human iPSCs were differentiated to NSC under chemically defined and xeno-free conditions ^4^. On day –2, 40,000 iPSCs per well were plated on vitronectin-coated 12-well plates in StemMACS iPS-Brew XF medium, supplemented with 2 μM Thiazovivin. The next day, the medium was replaced with fresh StemMACS medium. On day 0, neural differentiation was induced by changing the medium to neural induction medium (50% DMEM/F12, 50% Neurobasal medium, 1 × N2-supplement, 1 × B27-supplement, 1 × Glutamax, 10 ng/ml hLIF, 4 µM CHIR99021, 3 µM SB431542, 2 µM Dorsomorphin, 0.1 µM Compound E). The following day medium was changed. On day 2, the medium was switched to Neural Induction Medium 2 (50% DMEM/F12, 50% Neurobasal medium, 1 × N2-supplement, 1 × B27-supplement, 1 × Glutamax, 10 ng/ml hLIF, 4 µM CHIR99021, 3 µM SB431542, 0.1 µM Compound E) for four additional days and changed every day. On day 6, cells were transferred to pLO/L521-coated plates in Neural Stem Cell Maintenance Medium (50% DMEM/F12, 50% Neurobasal medium, 1 × N2-supplement, 1 × B27-supplement, 1 × Glutamax, 10 ng/ml hLIF, 4 µM CHIR99021, 3 µM SB431542) with daily medium changed, and cell passaging at 80–100% confluency. 2 μM Thiazovivin was added after splitting for the first six passages. From passage two onward, medium was supplemented with 5 ng/ml FGF2. NSC generated with this protocol show stable expression of various stem cell markers over at least 15 passages ^4^. At passage 5, cells underwent RNA switch-based purification as previously described ^4,21^. In brief, template DNA for *in vitro* transcription of mRNA was amplified by PCR from a vector encoding for either Barnase, Barstar, or puromycin using appropriate primers with the T7 promoter and poly(A) tail. The template of miR-302a-5p responsive OFF switch (Barstar) was amplified from the same vector by PCR using the primer containing miRNA anti-sense sequence at 5’UTR. For the miR-302a-5p responsive ON switch (Barnase), the open reading frame, poly(A) tail, and miRNA anti-sense sequence were amplified from the same vector by the first PCR. Then, the PCR product and extra sequence were fused in this order by a second PCR. All template DNAs were purified using the Min-Elute PCR Purification Kit. The RNAs were transcribed for 6h at 37 °C using MEGAScript T7 Transcription Kit using 1-Methylpseudouridine-5’-Triphosphate and Anti Reverse Cap Analog, ARCA. The transcribed RNA was treated with Turbo DNase I and antarctic phosphatase to remove template DNA and purified by RNeasy MinElute Cleanup Kit (QIAGEN). The concentration was determined by Qubit microRNA Assay Kit. One day before transfection, cells were passaged as detailed above and seeded onto PLO/L521 coated plates. The following day, mRNA was transfected using Lipofectamin RNAiMAX Transfection Reagent. The transfection complex was added to the cells in a dropwise manner and the plate was agitated before being placed into the incubator, followed by a medium change after 4h. To eliminate residual iPSCs and non-transfected cells, puromycin selection was performed (2 µg/ml, 24 h), leveraging Barnase-mediated degradation of the puromycin resistance mRNA in iPSCs.

### scFv construct generation

ScFv constructs were ordered from geneart (thermofisher) and cloned into pUKCCL vector (see Vector maps in **Suppl Figure 2**).

### qPCR

RNA extraction was carried out using Quiagen RNeasy kit according to the manufacturer’s recommendations. qPCR was performed using SYBR green kit (iTaq Universal SYBR Green Supermix from Biorad) containing 0.5 µM of each primer with the following cycling conditions (hold stage: 95 °C, 10 min, 1 cycle; PCR stage (95 °C, 15 s, 60 °C 1 min; 95 °C 15 s, 40 cycles; Melting curve (95 °C, 15 s, 60 °C, 1 min).

### Flow cytometry for integrin and protein L binding

NSC *were* washed in PBS, centrifuged at 300g for 5 min, and resuspend in 1 ml FACS buffer. 0.5 mio cells/well were plated on ABgene V-bottom 96-well Storage Plate (Thermo). Cells were centrifuged again for 5min at 400g and incubated with 100ul of 10 µg/ml his-tagged αvβ3 integrin or protein L in FACS Buffer for 30 min at at 4°C. After two centrifugation/wash steps in FACS buffer, cells were resuspended in 100 ul of anti his-tag PE + live/dead staining in FACS buffer and incubated for 30min at 4°C in the dark. After two centrifugation/wash steps in FACS buffer cells were fixed in 100 ul 1 % PFA and incubated for 20min at RT in the dark. Cells were again washed two times and acquired using a LSR Fortessa (BD Bioscience) and the data were analyzed by FlowJo software (Tree Star).

### Cell-based ELISA (cELISA)

96 well plates (NUNC-Immunoplate) were coated with 8 µg/ml αvβ3 integrin or 1 µg/ml protein L in PBS overnight at 4°C. Plates were washed, with 0.005 % Tween in PBS and blocked in blocking/dilution buffer (2% BSA, 0.2%Tween20, PBS) for 1.5 h at RT, slightly shaking. NSC were centrifuged at 300g for 5 min, resuspended in dilution buffer. Plates were washed, and cells were plated at 0.5 × 10^6^ cells/well in a serial dilution (0.5, 0.125, 0.062 × 10^6^, and 0 cells/well) and incubated while slightly shaking at at RT for 1.5h. After washing, for cell detection, 1 µM Hoechst was added and incubated for 20 min and Hoechst fluorescence was measured using fluorescence plate reader at 350 nm excitation and 461 nm emission wavelength (Tecan Infinite m1000 pro).

### In vitro luciferase assay

Cells were seeded on 24-well plates and incubated with 150 µg/ml luciferin for 10 min at 37 °C. Bioluminescence was measured with Tecan M1000 pro.

### Photothrombotic stroke induction

Photothrombotic stroke was induced as previously described ^53,56,57^. Anesthesia was initiated with 4% isoflurane in oxygen and maintained at 1–2% via a custom-made face mask. Once a deep anesthetic state was confirmed (breathing rate ∼50 breaths/min and absence of toe pinch reflex), mice were secured in a stereotactic frame (David Kopf Instruments). Body temperature was maintained at 36–37 °C with a heating pad, and eyes were protected with lubricant (Vitamin A, Bausch & Lomb). The head was shaved, and topical Emla™ cream (5% lidocaine/prilocaine) was applied to the scalp and ears before positioning in ear bars.

A ∼1 cm midline scalp incision was made to expose bregma and lambda. The skull surface was cleaned, and the target region for stroke induction (−2.5 to +2.5 mm mediolateral, 0 to +3 mm anteroposterior from bregma) was identified under a surgical microscope (Olympus SZ61) and marked. Rose Bengal (15 mg/ml in 0.9% NaCl) was injected intraperitoneally (10 μl/g body weight) 5 min before illumination. A cold light source (Olympus KL1500LCD, 150 W, 3000 K) was applied to the marked region (4 × 3 mm area) for 10 min.

Once illumination ended, the mouse was placed into an empty cage. 30 minutes post-surgery, the mouse’s skull was imaged using LDI to confirm stroke induction. After stroke verification, the incision was sutured using a Braun surgical suture kit. Betadine® was applied to the suture, and the mouse was returned to an empty cage, followed by standard post-op care.

### NSC transplantation

Transplantation of NSC was performed as described before ^27^. Operation procedures remained the same with photothrombotic stroke surgery up to the illumination step. After locating bregma, the stereotactic frame was adjusted to the injection coordinates [AP: +0.5, ML: +1.5, DV: –0.6 mm relative to bregma]. A ∼0.8 mm burr hole was drilled until the cortical bone was penetrated. A 10 μl Hamilton syringe with a 30G needle containing 2.5 μl of cell suspension (10⁵ cells/μl) was mounted on the stereotactic arm and positioned above the target site. The needle was lowered slowly to the calculated depth, with an additional 0.05 mm penetration to create a pocket for the graft. The needle was then retracted to the intended depth and cells were injected into the parenchyma at a constant rate of 2 nl/s. Following injection, the needle was kept in place for 5 min to allow the suspension to settle before gradual withdrawal. The cavity was sealed with Histoacryl®, and once hardened, the incision was sutured. Animals were returned to an empty cage and received standard post-operative care.

### Bioluminescent imaging (BLI)

Animals were imaged using the IVIS® Lumina III In Vivo Imaging System. For optimal signal detection, the entire head was shaved with an electric razor. Mice received an intraperitoneal injection of diluted D-luciferin in PBS (300 mg/kg body weight, sterile-filtered through a 0.22 μm syringe filter). Images were acquired 10, 15, 20, and 25 min after substrate administration. Data were analyzed using Living Image software (v4.7.3). A region of interest (ROI; 1.8 cm × 1.8 cm) was placed over the bioluminescent signal between the ears and nose. Two additional ROIs were defined: one at the posterior body to quantify background, and one at a randomly selected area near the animal to assess image noise. Signal intensity was measured as total photon flux (photons/s). Data were recorded in Microsoft Excel and further processed in R/RStudio.

### Laser Doppler imaging (LDI)

The surgical set-up was cleaned, and mice were placed in a stereotactic frame. Shortly after stroke induction the skull was exposed through a midline skin incision. Mice were subjected to LDI (*Moors Instruments, MOORLDI2-IR)*. LDI data were exported, quantified as flux within the defined ROI using Fiji (ImageJ), and further analyzed in RStudio.

### Tissue processing and immunohistochemistry

Animals were perfused transcardially with Ringer solution (0.9% NaCl). For RNA sequencing, brains were removed and immediately snap-frozen in liquid nitrogen. For immunohistochemistry, animals were perfused with Ringer solution followed by 4% paraformaldehyde (PFA). Brains were post-fixed in 4% PFA for 6 h, cut into 40 μm coronal sections using a Thermo Scientific HM 450 microtome, and rinsed in 0.1 M phosphate-buffered saline (PBS). Sections were incubated for 1 h at room temperature in 500 μl blocking buffer (5% donkey serum in PBS with 0.1% Triton® X-100), then incubated overnight at 4 °C with primary antibodies (**Suppl. Table 1**) on a Unimax 1010 shaker (∼90 rpm). The following day, sections were washed and incubated with appropriate secondary antibodies (**Suppl. Table 2**) for 2 h at room temperature, followed by counterstaining with DAPI (Sigma, 1:2000 in PBS). Sections were mounted on Superfrost Plus™ slides in 0.1 M PBS and coverslipped with Mowiol® mounting medium.

To visualize neurite outgrowth from the transplanted cell grafts, we performed 3,3′-Diaminobenzidine Tetrahydrochloride (DAB) staining using a human-specific neural cell adhesion molecule (hNCAM) in a separate cohort of animals at 28 days post-stroke (21 days post-transplantation), a timepoint chosen to capture later-stage axonal sprouting. Sections were treated with an antigen retrieval buffer at 70° for 30 min, followed by incubation in quenching solution for another 30 min. Sections were blocked using blocking buffer (BSA (5%), Glycin (1.5%), Triton X-100 (0.25%) and Casein (1%)) and subsequently incubated with anti-NCAM antibodies (1:10000, Abcam, #ab75813) at 4° overnight. The next day, sections were incubated with the secondary antibody (1:500, Vector Labs #BA-1000) for 2 hours at room temperature. Sections were then treated with the Avidin-Biotin-Complex (ABC) kit solution (Vector Laboratories, #PK-6100) according to the manufacturer’s protocol. The DAB staining was performed at a concentration of 0.5mg/ml DAB and 30% H_2_O_2_. Coronal sections were imaged on the Zeiss Axio Scan.Z1 slide scanner and processed using Fiji (ImageJ).

### Microscopy stroke area/volume quantification

Imaging of coronal brain sections was performed with a *Leica SP8 laser confocal microscope* with 10x and 63x objectives, and images were processed using *Fiji* (ImageJ) as previously described ^57^. For stroke analysis and graft size estimations, 40 μm coronal sections were stained with HuNu (graft) and GFAP (lesion size) and imaged on the Zeiss *Axio Scan.Z1* slide scanner and processed using *Fiji (ImageJ)*. The lesion area of each coronal section was measured by manually drawing a polygonal area around the stroke region as closely as possible. Brain sections were identified according to stereotaxic coordinates (distance relative to Bregma) using a reference mouse brain atlas ^58^. A custom-made RScript was used to re-scale the image and enhance contrast. Additional parameters (height, width) were recorded for graft size analysis. To assess lesion-and graft size volume, length, width, and area were measured for each brain section and ‘stacked together’ to create a 3-D model. Lesion- and Graft volume was approximated as an elliptical oblique cone:

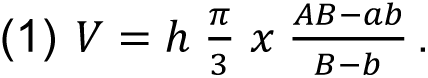

### Lesion coverage analysis

To quantify graft coverage of the stroke lesion, 40 μm coronal brain sections were stained with HuNu to identify grafted cells and GFAP to delineate the lesion area. Sections were scanned on the Zeiss Axio Scan.Z1 slide scanner and processed using Fiji (ImageJ). Graft and lesion regions were defined as regions of interest and analyzed using the ROI Manager. For each section, an area fraction was calculated as the ratio of graft area to stroke area. In parallel, an overlap coefficient was determined as a similarity measure between graft and lesion regions, defined as the area of intersection divided by the size of the smaller region. The lesion coverage index was then calculated by multiplying the area fraction by the overlap coefficient, resulting in a value between 0 and 1, with 1 indicating complete lesion coverage by the graft and 0 indicating no overlap.

### Histological quantification of vasculature, microglia and astrocytes

All analysis steps were performed in a peri-infarct region, referred to as ischemic border zone (IBZ) adjacent to the stroke core that extended up to 300 μm. Post-ischemic vascular repair was assessed using the software FIJI (ImageJ, version 2.1.0/1.53c) and previously established script ^59,60^ that allows to automatically calculate (1) vascular density, (2) number of branches and junctions and (3) the length of blood vessels^61^. The intensity of the glial scar and inflammation signal was determined by analyzing three brain sections per animal, which were immunostained for GFAP and Iba1. Images were converted to 8-bit format and binarized by applying thresholds derived from mean gray values of ROIs in the intact contralateral cortex. The total area corresponding to reactive gliosis or inflammation within and adjacent to the ischemic lesion was subsequently measured.

Morphology of Iba1^+^ cells was performed using the skeletal analysis plugin and/or the Sholl Analysis plugin in Fiji (ImageJ). Z-stack images (acquired in 63x) were projected using a maximum intensity projection. The brightness/contrast of the z-stack was then adjusted to best visualize the branches of the Iba^+^ microglia. Images were converted to grayscale, then set to 8-bit and thresholded using mean gray values from ROIs in the contralateral, unaffected cortex to generate binary masks. Circularity was quantified with the Analyze Particles plugin in ImageJ, where a value of 1.0 corresponds to a perfect circle and values near 0.0 indicate highly elongated shapes.

For calculating the branching and the ramification index, images were skeletonized with the plugin Analyze Skeleton^62^ and followed by the Sholl analysis plugin ^63^. A radius was drawn from the middle of the cell body to the end of the most distant branch of the cell to set the lower and upper limit for concentric circle placement. The first circle was set close to the edge of the cell body to ensure the cell body was not counted as an intercept on the circle. The distance between the circles was set as 2 µm for all analyzed cells.

### Human RNA-seq analysis

Uniformly processed gene-level read counts for GSE56267 (7 ischemic stroke cortex, 6 healthy cortex; median time after stroke: 3 years; Illumina HiSeq 2000) were obtained from the NCBI GEO RNA-seq Experiments count repository (GRCh38.p13, HISAT2 alignment, Subread featureCounts, NCBI Annotation Release 109). Differential expression analysis was performed in R using DESeq2 (v1.44) with design formula ∼condition (stroke vs. healthy). Genes with fewer than 10 counts in at least 6 samples were excluded. Integrin family subunit expression was visualized as log_2_fold change centered to the healthy group mean.

### Statistical analysis

Statistical analyses were performed using RStudio (Version 4.0.4). Sample sizes were determined to ensure adequate statistical power based on prior studies from our group and relevant literature ^28,29,38,64,65^. Data distribution was assessed using the Shapiro–Wilk test. Normally distributed data were analyzed using unpaired, two-sided t-tests for comparisons between two cell lines. For comparisons involving multiple groups or time points (e.g., longitudinal cell survival data), differences were evaluated using ANOVA followed by post-hoc analysis of estimated marginal means. Non-normally distributed data were analyzed using an unpaired Wilcoxon–Mann–Whitney test. Data are presented as mean ± SD; or mean ± SEM, statistical significance was defined as *p < 0.05, **p < 0.01, and ***p < 0.001.

## Data availability

All source data are available for reproducibility purposes from the corresponding author upon request. Source data are provided with this paper. Human stroke RNAseq data were obtained from GSE56267.

## Conflict of interest statement

The authors declare that the research was conducted in the absence of any commercial or financial relationships that could be construed as a potential conflict of interest.

## Acknowledgements

This work is supported by the funding from the Mäxi Foundation, Swiss 3R Competence Center (OC-2020-002), Swiss National Science Foundation (CRSK-3_195902) and (PZ00P3_216225), USC Dean’s Pilot Funding Award (000092), and the Neuroscience Center Zurich.

## Author contributions

R.R., R.Z.W., and C.T. designed the study. R.R., R.Z.W., B.A.B., and N.H.R. conducted and analyzed in vivo experiments. C.B., K.J.Z., D.U., D.M., and U.K. planned, conducted, and analyzed in vitro experiments. R.R., R.Z.W., B.A.B., M.Z., and C.T. generated figures. M.G. provided an iPS cell line. H.S. provided the RNA switch system. R.R., R.M.N., S.P.H., and C.T. supervised the study. R.R., R.Z.W., P.H., and C.T. wrote the manuscript with input from all authors. All authors read and approved the final manuscript.

## Notes

### Competing Interest Statement

The authors have declared no competing interest.

