## Supplemental Figures for "Targeting integrin αvβ3 by chimeric antigen receptor neural stem cell (CAR-NSC) therapy for stroke"

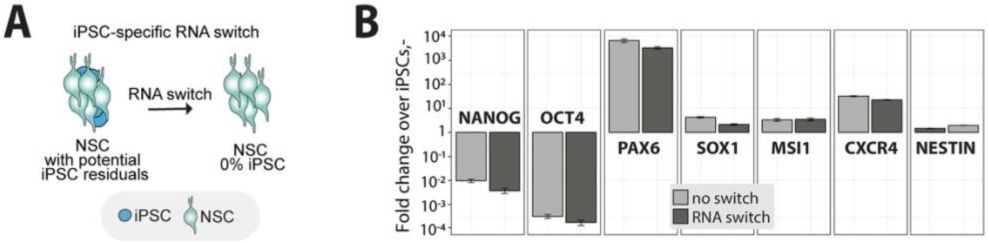


**Suppl. Figure 1: Gene expression of hiPSC-derived NSC before and after RNA switch.** qPCR showing fold change over iPSCs of iPSC marker NANOG and OCT4 and NPC marker PAX6, SOX1, MSI1, CXCR4 and Nestin in NSC without (light gray) and with RNA switch (dark grey).

A

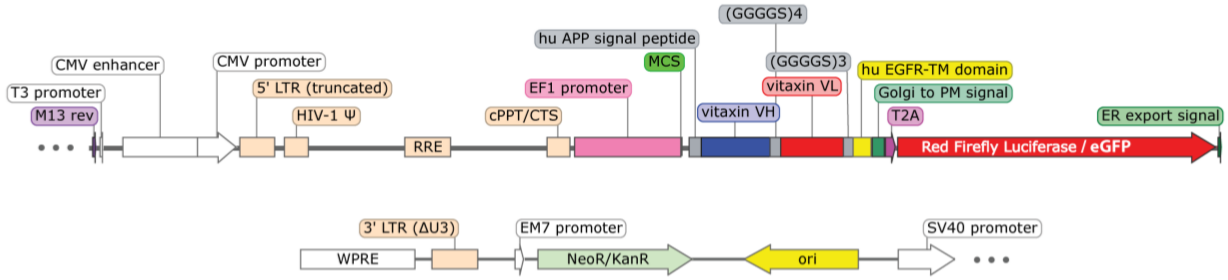


B

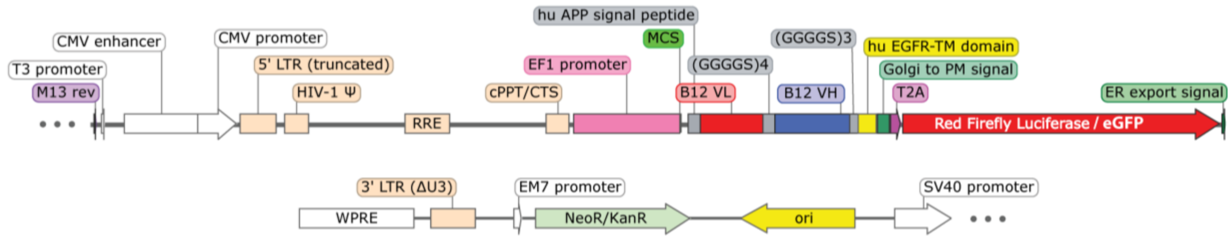


**Suppl. Figure 2: Map of scFv expression vectors**. (A) Schematic representation of pUKCCL plasmids for the expression of αvβ3-scFv and (B) the B12 control scFV. VL: variable light chain; VH: variable heavy chain.

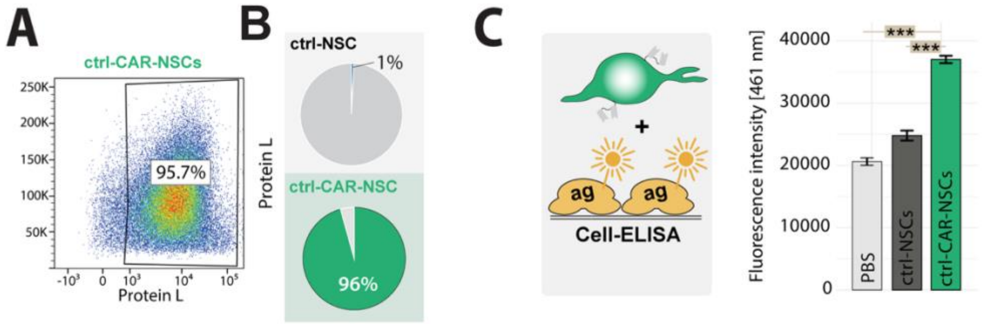


**Suppl. Figure 3: Control-CAR-NSC binding to Protein L.** (A) Representative flow cytometry dot plot of control-CAR-NSC, incubated with recombinant his-tagged protein L. (B) Quantification of flow cytometry analyses. The percentage of non-transduced NSC and control-CAR-NSC that bound to his-tagged protein L is shown. (C) Cell-based ELISA showing Hoechst fluorescence intensity of cells bound to protein L-coated plates. Each bar represents the mean relative fluorescence signal of two wells seeded with 500000 cells (ctrl-NSC and ctrl-CAR-NSC) or PBS. Statistical significance was assessed using Tukey's HSD. *$p < 0.05$, **$p < 0.01$, ***$p < 0.001$.

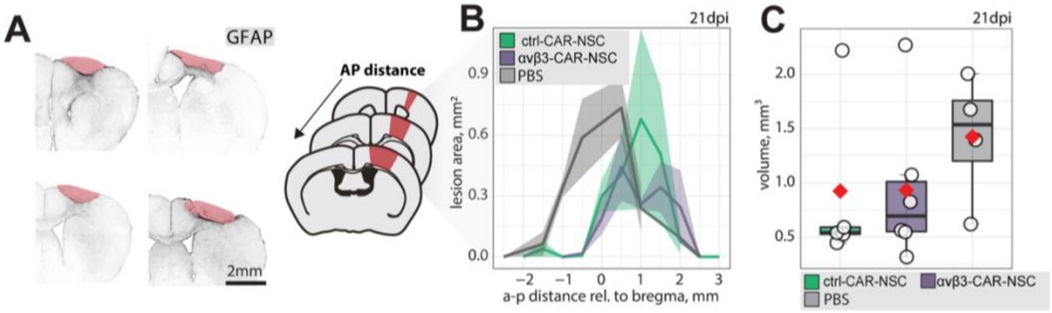


**Suppl. Figure 4: Stroke lesion size assessment.** (A) Representative GFAP staining outlining the ischemic lesion along the A-P axis. (B) Lesion area plotted against the anterior-posterior (a–p) distance relative to bregma (mm). (C) Quantification of lesion volume across groups. Box plots show the median (center line), 25th to 75th percentiles (bounds of the box), and minimum to maximum values (whiskers). Red dot indicate the mean value per group. Each dot represents one biological replicate (individual animal). Line graphs are plotted as mean ± sem. Statistical differences were assessed using one-way ANOVA followed by Tukey's HSD. $^{*}p < 0.05$, $^{**}p < 0.01$, $^{***}p < 0.001$.

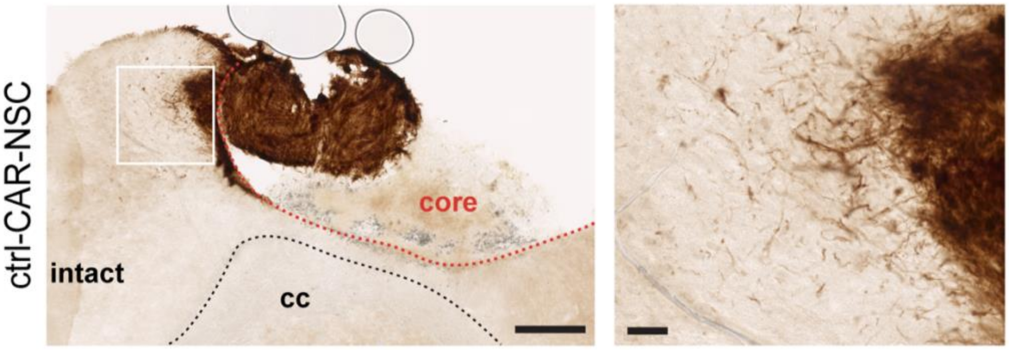


**Suppl. Figure 5: Representative human-specific NCAM (hNCAM) immunostaining showing neurite outgrowth from ctrl-CAR-NSC grafts in the peri-infarct cortex.** Left: overview with intact cortex, stroke core (core), and corpus callosum (cc). Right: magnified views of boxed region.

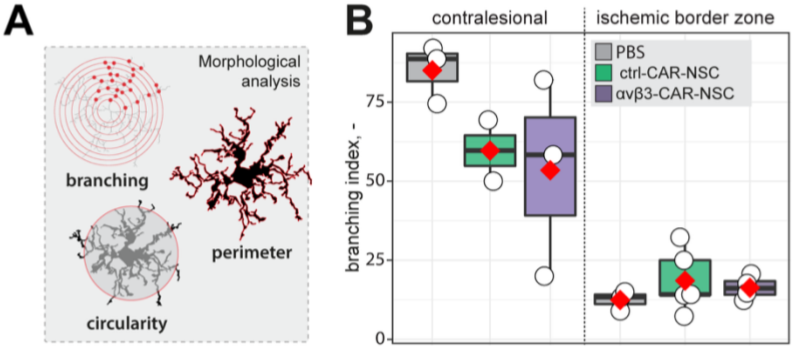


**Suppl. Figure 6. Microglial activation in PBS-, ctrl-CAR-NSC- and αvβ3-CAR-NSC-treated mice.** (A) Schematic of morphological parameters quantified: branching index (B) Quantification of branching index in contralesional cortex and ischemic border zone for PBS, ctrl-CAR-NSC, and αvβ3-CAR-NSC groups.
